# TurbOmics: a web-based platform for the analysis of metabolomics data using a multi-omics integrative approach

**DOI:** 10.1101/2025.05.09.653072

**Authors:** Rafael Barrero-Rodríguez, Jose Manuel Rodríguez, Thomas Naake, Wolfgang Huber, María Juárez-Fernández, Almudena R. Ramiro, Jesús Vázquez, Annalaura Mastrangelo, Alessia Ferrarini

## Abstract

In recent years, multi-omics integration has proven highly effective for the holistic characterization of biological systems, with the development of numerous bioinformatic platforms. However, these tools face limitations when incorporating metabolomics data, including absence of support for untargeted metabolomics annotation and dependence on predefined knowledge bases. Furthermore, the advanced algorithms required for multi-omics integration typically demand programming skills and statistical background, restricting their use to specialized users. To advance towards resolving these challenges, we present TurbOmics, a user-friendly web-based platform that enables researchers with diverse backgrounds to analyze metabolomics, proteomics, and transcriptomics data using advanced algorithms for multi-omics integration, while addressing key challenges associated with metabolomics data. Users can upload quantitative data and include additional information, such as metabolite identification or lipid classes, that streamline the interpretation of complex results. TurbOmics can be used sequentially with our previously published tool, TurboPutative, to simplify, reduce and prioritize the list of putative annotations from untargeted metabolomics datasets. Thanks to its flexible and interactive interface, researchers can perform exploratory data analysis and multi-omics integration using the Multi-Omics Factor Analysis, Pathway Integrative Analysis and Enrichment Analysis modules. We believe that TurbOmics will make multi-omics analysis more accessible to the research community. The platform is freely available at https://proteomics.cnic.es/TurboPutative/TurbOmicsApp.html.

GRAPHICAL ABSTRACT

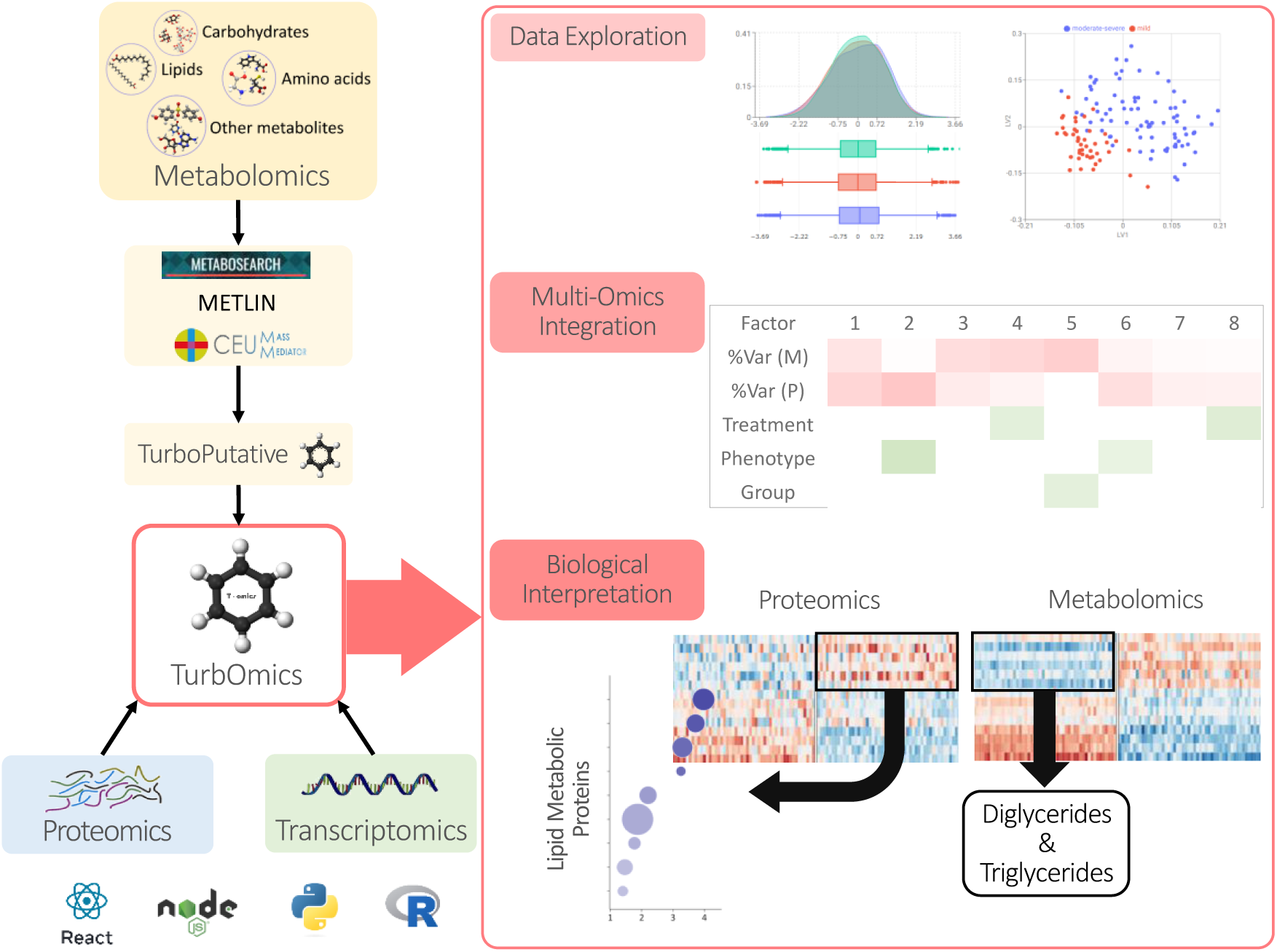

## INTRODUCTION

Multi-omics integration has undergone significant technological advancements in recent years both in next generation sequencing (1) and mass spectrometry (2, 3) that enable the simultaneous probing of multiple omics layers within a single study. The combined analysis of multiple omics data has proven to be highly effective for the molecular characterization of multifactorial diseases (4–6), the identification of novel disease biomarkers (7–9), and the advancement of personalized medicine (10–12). However, despite the availability of various bioinformatics tools designed for multi-omics integration, important limitations persist, particularly in incorporating results from emerging fields such as metabolomics (13, 14).

Metabolomics aims at measuring the metabolites present within a cell, tissue or organism at a specific time and under particular conditions (15, 16). Metabolomics strategies are traditionally divided into two approaches: targeted and untargeted. Targeted metabolomics measures a defined set of known metabolites, while untargeted metabolomics involves the comprehensive analysis of all analytes measurable in a sample (17). Both approaches commonly employ hyphenated techniques coupled to mass spectrometry (MS) (*e.g.*, liquid/gas chromatography coupled to MS), and, to a lesser extent, Nuclear Magnetic Resonance (NMR) spectroscopy to measure metabolites in biological samples (15, 16, 18). Processing and analyzing metabolomics data pose challenges that range from metabolite identification to biological interpretation at functional and metabolic pathway levels. To address these challenges, numerous platforms have been developed in recent years for data pre-processing and analysis, including MetaboAnalyst (19), XCMS Online (20) and Workflow4Metabolomics (21). In this context, we previously developed TurboPutative, a tool for streamlining data handling, classification and interpretation of untargeted metabolomics data (22). However, important limitations still persist in analyzing both targeted and untargeted metabolomics data particularly regarding its integration with other omics (14).

To address these challenges, several web applications for multi-layer omics analysis have emerged, facilitating omics data analysis and lowering the barrier to the use of complex machine learning algorithms and multivariate statistical models (23). However, these tools that are highly useful and versatile for the integration of genomics, transcriptomics and proteomics datasets, often fall short when analyzing metabolomics data within this multi-omics framework. For instance, knowledge-driven tools like PaintOmics (24), MergeOmics (25) and OmicsNet (26) map genes or metabolites of interests to known pathways but are constrained by the quality and coverage of their underlying databases. Conversely, data-driven tools such as 3Omics (27), MiBiOmics (28) or OmicsAnalyst (29), although more general and unbiased, do not support annotation of untargeted metabolomics features or the combination of researcher annotations with subsequent multivariate statistical analyses. Thus, there is an urgent need for a bioinformatic tool that 1) supports multi-omics analysis of both targeted and untargeted metabolomics data, addressing the annotation challenges posed by the latter, 2) facilitates the exploration of user-provided annotation and metabolite identifications to improve biological interpretation and coverage, and 3) provides a user-friendly interface offering interactive visualizations and data exploration capabilities.

To address these limitations, we introduce TurbOmics, a web-based platform for the comprehensive analysis and visualization of metabolomics data using a multi-omics integrative approach. The platform supports data from targeted-based experiments or those where the metabolite’s identification is available (i.e. NMR-or gas chromatography coupled to MS-based experiments). Furthermore, TurbOmics enables users to putatively annotate untargeted metabolomics features based on their mass-to-charge (m/z) ratio and incorporates additional user-provided information, such as metabolite names and lipid classes, streamlining the interpretation of complex results. Within TurbOmics, researchers can perform exploratory data analysis and multi-omics integration using the Multi-Omics Factor Analysis, Pathway Integrative Analysis and Enrichment Analysis modules. The platform integrates multivariate statistical models with functional pathways from knowledge bases such as KEGG (30) and Reactome (31), enabling the mapping of identified biomolecules and facilitating biological interpretation of the results. Users can explore and export data tables, visualize plots, and gain insight into the biological system under study. With its highly interactive interface, this website is free and open to all users with no login requirement.

## RESULTS

TurbOmics accepts transcriptomics, proteomics, and metabolomics data collected from the same set of biological samples. Metabolite identifications or putative annotations of untargeted metabolomics features, which are obtained using tools such as CMM (32), MetaboSearch (33), or METLIN (34) and further processed with TurboPutative, can be uploaded to the platform. After data pre-processing, users can investigate the results using the Exploratory Data Analysis section and perform multi-omics integration through the Multi-Omics Factor Analysis, Pathway Integrative Analysis and Enrichment Analysis modules (Figure 1).

**Figure 1.**
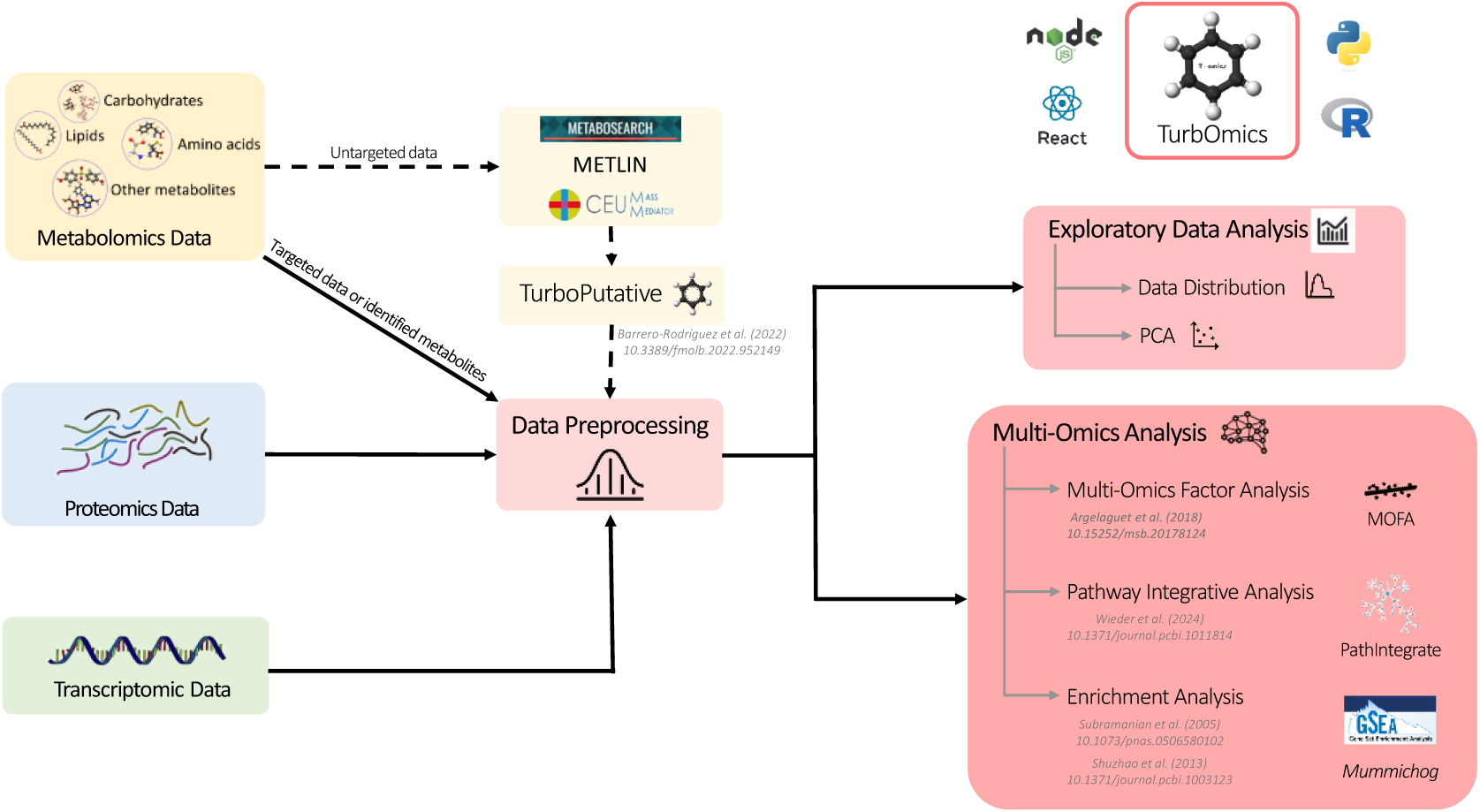
TurbOmics workflow.

### Data processing

#### Data upload

TurbOmics accepts data tables containing biomolecule abundances or quantifications generated from transcriptomics, proteomics, and metabolomics experiments. In order to perform exploratory and integrative analysis, users must upload a table containing metadata related to the biological samples (*e.g.*, treatment, sex, time). Optionally, additional omics metadata tables can be provided for each omics type to include relevant biomolecule information. For instance, users can include details regarding feature identification, putative annotations, lipid class, and biochemical pathways, which can enhance metabolomics analysis and improve biological interpretation (Supplementary Figure S1). All tables must be uploaded in TSV format. Alternatively, TurbOmics supports uploading data as an RDS file containing a MultiAssayExperiment object, simplifying the platform’s usability for the bioinformatics community familiar with Bioconductor (35).

#### Data preprocessing

Users can choose to apply variance stabilizing normalization, median normalization or logarithm transformation to their data. Additionally, biomolecule abundances and quantifications can be centered and scaled to improve the integration of different omics datasets. Biomolecules can also be filtered based on the presence of missing values, which can be imputed using various methods such as KNN, minimum, mean, or median.

#### Putative annotations

Putative annotations for untargeted metabolomics features can be uploaded to the platform in the metabolomics metadata table. Alternatively, users can obtain these annotations through TurbOmics, which integrates CMM and TurboPutative for this purpose. To do so, users must specify the columns containing the m/z ratio, retention time, and ionization mode for each feature. Additionally, the user can set the mass tolerance (ppm) and possible adducts present in the experimental data, according to the specific analytical method employed to analyze the samples. The user is required to provide the parameters used by the TurboPutative TPMetrics module to calculate a confidence score for the annotations, as previously described in (22). Briefly, TPMetrics applies a multi-criteria scoring algorithm where the user needs to define the possible adducts for each metabolite class and the timeframe for the pairwise correlation among annotations within a narrow retention time window. After execution, the user will be able to access the tables with the putative annotations through a link to TurboPutative. The content of the final TurboPutative table, TPFilter, which contains summarized information of the putative annotation, will be used in the subsequent multi-omics integrative analysis.

### Multi-omics integrative analysis

#### Exploratory Data Analysis

TurbOmics enables users to explore the distribution of biomolecule abundance or quantification and assess data quality for each individual omics dataset. The platform provides interactive density curves and boxplots, allowing data distribution visualization by sample sets defined by the user. In addition, users can explore distributions for specific biomolecule classes (*e.g.*, triglycerides, phospholipids, mitochondrial proteins). TurbOmics also provides dimensionality reduction techniques, which offer low-dimensional representations of high-dimensional omics data, revealing major trends and characteristics. The platform includes PCA for each omics dataset, allowing users to select among scatter plots of sample projections and analyze biomolecule loadings for each component. These functionalities facilitate the rapid detection of outliers, batch effects and relevant biomolecules for group comparisons.

#### Multi-Omics Factor Analysis

TurbOmics includes the MOFA statistical method for the unsupervised integration of quantitative data across multiple omics. MOFA identifies latent factors that capture the main sources of variation in the analyzed data, enabling the visualization of subgroups among samples as well as shared patterns across omics layers (36). TurbOmics performs univariate linear regression between the sample projections on each factor and the associated metadata, allowing for the rapid identification of biologically relevant factors. The platform offers an interactive interface for exploring the biomolecules that contribute the most to each factor, generating heatmaps and providing exportable tables with detailed information for data analysis. To support biological interpretation, TurbOmics performs ORA analysis on the proteins, transcripts and metabolites most associated with the factors of interest, offering biological insights into MOFA results.

#### Pathway Integrative Analysis

Pathway-based multi-omics integration is performed using both the Multi-View and Single-View PathIntegrate frameworks. PathIntegrate uses a supervised approach to model multi-omics data at the pathway level. Briefly, the Multi-View approach calculates pathway scores for each individual omics, providing interpretable insights both within and across omics. In contrast, the Single-View approach computes more comprehensive multi-omics pathway scores resulting from the integration of the quantitative information from multiple omics. Multi-omics analyses at pathway-level increase statistical power, facilitates biological interpretation, and enables the detection of multiple, weakly correlated signals that may not be detected individually (37). TurbOmics streamlines the use of the PathIntegrate framework by providing a user-friendly interface that simplifies execution and aids in result interpretation. Within TurbOmics, users can select either the Single-View or Multi-View supervised models, specify the comparison groups, and define the columns containing biomolecule identifiers. Upon execution, the browser presents a summary table with key statistical details of the model, alongside a scatter plot showing sample projections on the latent variables. Additionally, a table highlights the most relevant pathways with associated biomolecules. For the Multi-View model, separate pathway tables are generated for each omics dataset (Supplementary Figure S2). To facilitate biological interpretation, TurbOmics provides a heatmap visualization of biomolecule abundance or quantification values associated with the selected pathway.

#### Enrichment Analysis

TurbOmics offers a flexible and comprehensive section to perform enrichment analysis using a multi-omics approach. This section allows users to run the GSEA algorithm in parallel across transcriptomics, proteomics, and metabolomics data, facilitating the comparison of statistically significant pathways between these datasets. Users must specify the biomolecule identifier column and choose a metric for GSEA, such as LogFC or t-statistic. Additionally, PCA and MOFA loadings can be selected, enabling deeper analysis of the components and factors calculated in earlier sections. For untargeted metabolomics data, enrichment analysis can be performed using the mummichog library. Mummichog model has shown improved functional enrichment using m/z ratios and retention times. Specifically, the model firstly maps detected features onto potential metabolites in a network and then searches for local enrichments that reflect true biological activity. The results are displayed in an interactive and exportable table that includes the biomolecules responsible for the enrichment in each category. TurbOmics also supports custom enrichment analysis, allowing users to define and explore their own categories. This capability is particularly useful in untargeted metabolomics, where users can explore the abundance or quantitative profiles of specific lipid classes or metabolic pathways of interest that are not included in standard databases.

### Application of TurbOmics to untargeted metabolomics data in atherosclerosis

To illustrate the ability of TurbOmics in analyzing untargeted metabolomics data with a multi-omics approach, we have selected a previously published collaboration from our group (38). In this study a single cell analysis of the antibody repertoire in atherosclerosis identified A12, an antibody that binds aldehyde dehydrogenase 4 family member A1 (ALDH4A1) mitochondrial enzyme. Further, infusion of the A12 autoantibody delayed plaque formation and reduced circulating free cholesterol and LDL levels in an atherosclerosis mouse model (Ldlr-/-HFD). Here, we re-analyzed the liver proteomics and untargeted lipidomics data collected to study the effects of serial intravenous injections of the A12 antibody compared to mice injected with B1-8 control antibody or PBS. Additionally, we incorporated new RNA-seq data from the same samples into the analysis.

The transcriptomics, proteomics and metabolomics datasets contained 11, 999 genes, 4, 114 proteins and 2, 935 features (2, 226 in positive mode and 709 in negative mode), respectively. The CMM run yielded 132, 815 annotations (116, 867 in positive mode and 15, 948 in negative mode), which TurboPutative reduced to 3, 028 prioritized entries in the TPFilter tables (2, 333 in positive mode and 695 in negative mode), using the multi-criteria scoring algorithm TPMetrics described above.

In the Exploratory Data Analysis section, we confirmed high data quality across all omics datasets. PCA performed on each omics dataset revealed axes of variation associated with antibody treatment (Supplementary Figure S3A-C). Notably, the second principal component in the metabolomics dataset was closely associated with the A12 antibody treatment (Supplementary Figure S3D), with triglyceride features showing the highest positive loadings (Supplementary Table S1.1). A density plot analysis of triglyceride features annotated in LipidMaps also indicated a slight decrease in the A12-treated group (Supplementary Figure S3E).

Multi-omics integration was performed using Multi-Omics Factor Analysis section. The MOFA model identified seven factors, explaining 75.4% of the variability in transcriptomics, 55.6% in metabolomics and 61.2% in proteomics (Figure 2A). Factor-treatment association analysis suggested that factors 4 and 6 were associated with A12 treatment (Figures 2A and 2B).

**Figure 2.**
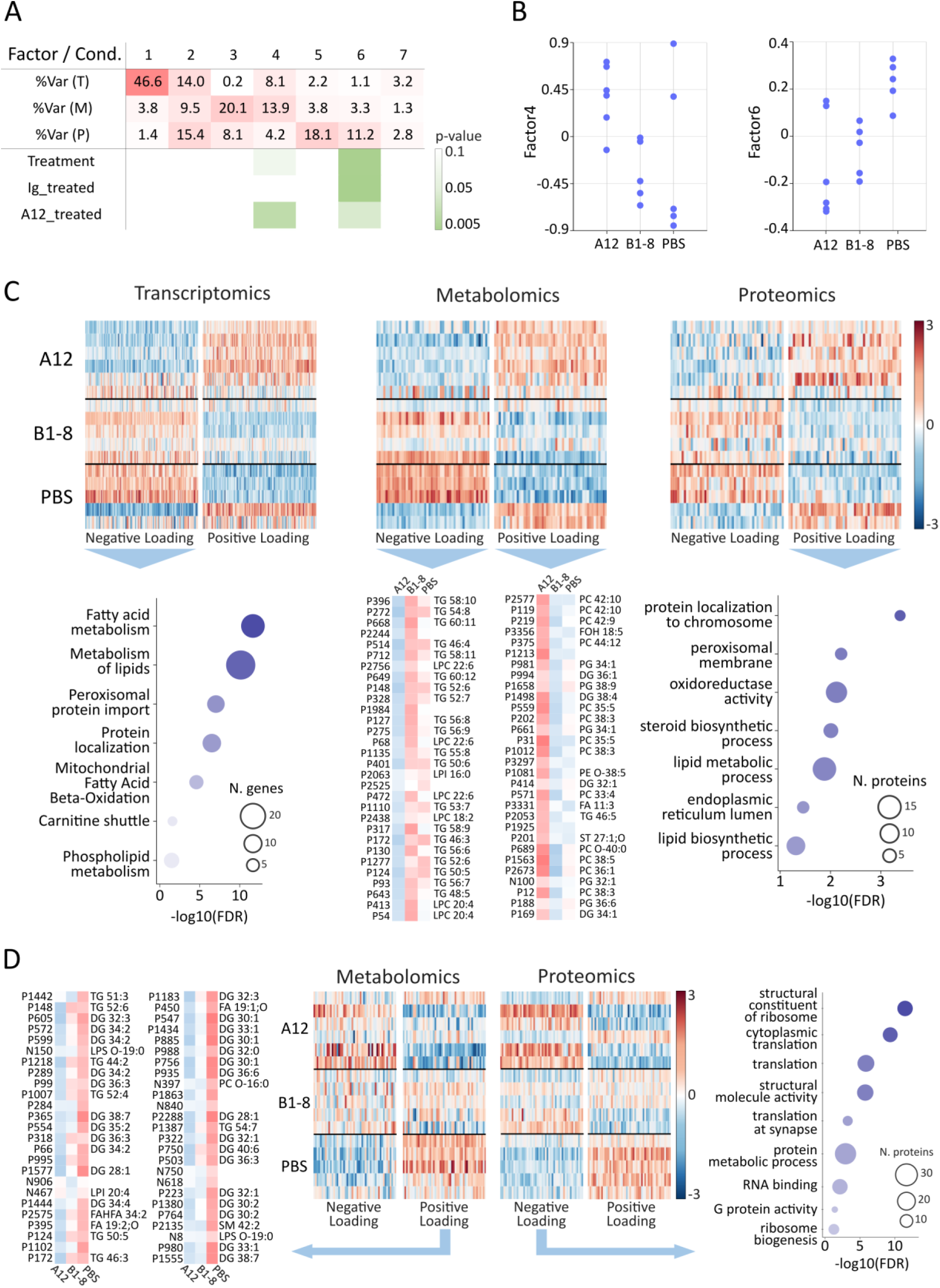
Multi-omics analysis of the atherosclerosis study using TurbOmics. (A) The table summarizes the MOFA factors, their percentage of explained variability in each omic layer, and the p-values for their association with the treatment groups. Three group comparisons are presented: *Treatment* (A12 vs. B1-8 vs. PBS), *Ig_treated* (A12 and B1-8 vs. PBS), and *A12_treated* (A12 vs. B1-8 and PBS). (B) Scatter plots display the distribution of Factor 4 (left) and Factor 6 (right) scores across treatment groups. The A12, B1-8, and PBS groups correspond to mice treated with A12, B1-8, and PBS, respectively, with the latter two serving as control groups. (C) The top panel presents heatmaps of the most associated elements with Factor 4 for each omic layer, including transcriptomics (120 genes per heatmap), metabolomics (50 metabolomics features per heatmap), and proteomics (50 proteins per heatmap), categorized separately by positive and negative loadings. The bottom left bubble plot shows enriched functional categories (ORA) among the 120 genes with the most negative loadings. The bottom middle panel contains tables listing the annotations of the 30 metabolomics features with the most negative loadings on the left and the 30 with the most positive loadings on the right. The bottom right bubble plot displays enriched functional categories (ORA) among the 50 proteins with the most positive loadings. (D) The top panel presents heatmaps of the most associated elements with Factor 6 for metabolomics (50 metabolomics features per heatmap) and proteomics (50 proteins per heatmap), categorized by positive and negative loadings. The bottom left table provides the annotations of the 50 metabolomics features with the most positive loadings, while the bottom right bubble plot shows enriched functional categories (ORA) among the 50 proteins with the most negative loadings.

Factor 4, though with varying intensity, was linked to metabolomics (metabolomics variance = 13.9%), transcriptomics (14.0%) and proteomics (4.2%). Genes, proteins and metabolomics features most associated with this factor exhibited similar abundance patterns, high correlation, and differential expression in A12 group (Figure 2C). Specifically, among the 50 metabolomics features with the highest negative loadings, which appeared decreased in A12 group with respect to controls, 30 were identified as triglycerides, and 8 as lysophosphatidylcholines (LPC), containing polyunsaturated fatty acids, including arachidonic and docosahexaenoic acid (Figure 2C, middle panel; Supplementary Table S1.2). Conversely, metabolomics features with positive loadings were enriched in phosphatidylcholines (PC), which were increased in the A12 group (Figure 2C, middle panel; Supplementary Table S1.3). Notably, previous studies have highlighted that serum PC and LPC profiles are altered in atherosclerotic patients (39). LPC is claimed to be an important component of Ox-LDL and has been detected in atherosclerotic lesions. However, its role in atherosclerosis remains complex, as it can exert both pro- and anti-atherogenic roles depending on the arterial cell type and the inflammatory context (40). In addition, certain circulating PC species have been reported to inversely correlate with the risk of myocardial infarction (41, 42), and PC has also been described to exhibit anti-inflammatory properties (43–45). Consistently, overrepresentation analysis (ORA) of the top 120 transcripts with the most negative loadings (decreased in the A12 group) showed enrichment in lipid and fatty acid metabolism categories (Figure 2C, left panel; Supplementary Table S1.4). Similarly, ORA on the top 50 proteins with the highest positive loadings revealed enrichment in lipid metabolism pathways (Figure 2C, right panel; Supplementary Table S1.5).

Factor 6 was highly associated with proteomics data (variance explained = 11.2%) and, to a lesser extent, with metabolomics (3.3%). Among the 50 metabolomics features with the highest positive loadings, 27 were diglycerides, and 7 were triglycerides, all of which were decreased in the A12-treated group (Figure 2D, left panel; Supplementary Table S1.6). Additionally, 30 out of the 50 proteins with the most negative loadings were associated with metabolic processes and protein synthesis, and were increased in A12 group (Figure 2D, right panel; Supplementary Table S1.7).

Overall, TurbOmics provided results consistent with the original study, identifying significant alterations in arachidonic acid metabolism, reductions in triglyceride levels, and changes in the expression of genes linked to lipid and amino acid metabolism. Importantly, these findings were obtained through an efficient and structured analysis workflow leveraging various TurbOmics modules. The application of the MOFA model and the interactive exploration of the generated results not only highlighted the role of PCs in atherosclerosis but also revealed additional biological insights that were previously undetected, such as the correlated alterations in PCs and LPCs in response to A12 treatment. Additionally, our analysis identified 37 decreased metabolomics features annotated as triglycerides and 27 as diglycerides, compared to the original study’s findings of 13 and 3, respectively. This difference is due to the original study’s approach of filtering for features showing statistically significant changes after multiple testing correction, which can lead to information loss and limit interpretative depth in multi-omics studies. By obtaining putative annotations for all metabolomics features and applying multivariate statistical models without pre-filtering, TurbOmics enables MOFA factors to capture complex biomolecular variations across datasets. This strategy mitigates noise and enhances the detection sensitivity of biomolecules with distinctive abundance patterns, ultimately providing a more comprehensive dataset for biological interpretation.

### Application of TurbOmics to targeted metabolomics data in a COVID-19 study

To demonstrate TurbOmics’ capability in analyzing targeted metabolomics data with a multi-omics approach, we have selected a COVID-19 study where plasma multi-omics profiles and circulating immune cell classes are explored by targeted metabolomics and proteomics approaches, in a large cohort of patients with varying infection severities [data sourced from Table S1 in (46)]. Patients were grouped based on disease severity, as quantified by the World Health Organization (WHO) ordinal scale (WOS), in mild (WOS = 1–2, n=49), moderate (WOS = 3–4, n=60), and severe (WOS = 5–7, n=30). Additionally, a cohort of healthy donors provided plasma samples for metabolomics (n=133) and/or proteomics (n=124) analyses, with an overlap of 18 individuals.

Quantitative plasma metabolomics and proteomics data were uploaded to TurbOmics along with metadata tables describing the sample groups. Additional metadata tables were provided for the metabolomics and proteomics data. For the metabolomics data, the table included metabolite names and corresponding ChEBI identifiers when available. For the proteomics data, the table contained protein names, their associated coding genes, and UniProt accession numbers. After preprocessing, the metabolomics dataset comprised 750 metabolites, while the proteomics dataset included 454 proteins.

PCA of metabolomics and proteomics data reveals clear distinctions between patients based on disease severity (Figure 3A). Of the 50 metabolites with the highest PC1 loadings, 24 are amino acid derivatives (Supplementary Table 2.1 and 2.2), with their quantitative values progressively increasing along disease severity (Figure 3B, left panel). In contrast, 24 metabolites with the most negative PC1 loadings are lipids (Supplementary Table 2.1 and 2.3), which exhibit a marked decrease in the moderate and severe groups (Figure 3B, right panel). Within the proteomics dataset, PC1 separates healthy individuals from patients, while PC2 differentiates patients based on disease severity. Proteins most strongly associated with PC1 include CD40 and CD274 (Supplementary Table 2.4), both of which have been previously identified as relevant in the context of COVID-19 (47, 48).

**Figure 3.**
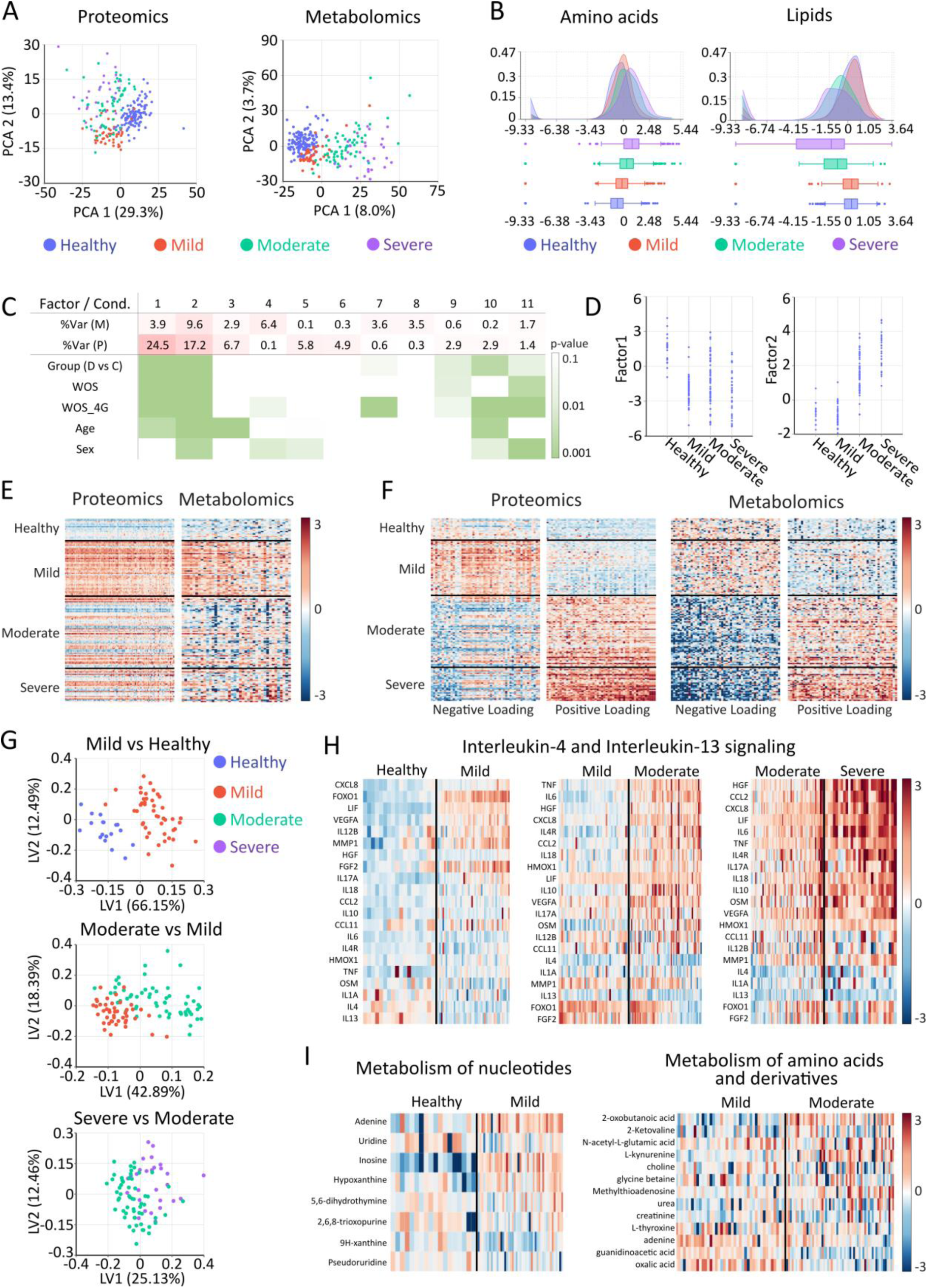
Multi-omics analysis of the COVID-19 study using TurbOmics. (A) Scatter plots showing the scores of Principal Components (PCs) 1 and 2 obtained from proteomics (left) and metabolomics (right) data. (B) Density and box plots illustrating the quantitative distribution of amino acids and derivatives among the 50 metabolites with the highest positive loadings on PC1 (left), and lipids among the 50 metabolites with the highest negative loadings on PC1 (right). (C) Table summarizing MOFA factors, their percentage of explained variability in each omic layer, and the p-values for their associations with patient groups. Abbreviations: D, disease; C, controls; WOS, disease severity (scale from 0 to 7); WOS_4G, groups studied (i.e. controls (WOS = 0), mild (WOS = 1–2), moderate (WOS = 3–4), and severe (WOS = 5–7)). (D) Scatter plots displaying the scores of Factor 1 (left) and Factor 2 (right) as a function of disease severity. (E) Heatmap displaying the quantitative values of the top 100 proteins most strongly associated with Factor 1, along with the 40 metabolites exhibiting the highest negative loadings within the same factor. (F) Heatmap depicting the 50 proteins and metabolites most strongly associated with Factor 2, categorized by positive and negative loadings. (G) Scatter plot of MB-PLS superscores for latent variables 1 and 2 derived from contrasts between mild vs. healthy, moderate vs. mild, and severe vs. moderate groups, as analyzed in the Pathway Integrative Analysis module. (H) Heatmaps of the quantitative values for proteins belonging to the category *Interleukin-4 and Interleukin-13 signaling* (R-HSA-6785807) across the different contrasts (top). Heatmaps of metabolite quantitative values for the category *Metabolism of nucleotides* (R-HSA-15869) in mild vs. healthy, and for the category *Metabolism of amino acids and derivatives* (R-HSA-71291) in moderate vs. mild (bottom).

Multi-omics integration using the Multi-Omics Factor Analysis module allowed the unsupervised identification of two complementary axes of variability highly associated with disease severity (Figure 3C). Factor 1 (proteomics variance = 24.5%; metabolomics variance = 3.9%) was found primarily associated with proteomics and distinguished healthy individuals from diseased groups (Figure 3D, left panel). In contrast, Factor 2 (proteomics variance = 17.2%; metabolomics variance = 9.6%) was associated with both omics layers and differentiated moderate-severe from healthy-mild groups (Figure 3D, right panel). Proteins with the highest weight in Factor 1 (Supplementary Table 2.5) are characterized by a notable increase in COVID-19 patients, especially in the mild group (Figure 3E, left panel). These include proteins involved in the NF-kappa B signaling pathway, such as CD40LG, IRAK1, IKBKG, or PRKCQ, as well as apoptotic proteins like AIFM1, TRAF2, or BIRC2, which align with findings from previous studies (49–51). Metabolites associated with this factor, though to a lesser extent, also exhibit a notable increase in mild COVID-19 patients (Figure 3E, right panel), including several sphingolipids like sphingosine 1-phosphate (Supplementary Table 2.6), which has been previously linked to COVID-19 severity (52, 53). Proteins and metabolites associated with Factor 2 can be categorized into two groups based on whether their levels increase or decrease in the moderate-severe patient groups (Figure 3F). The proteins increased in the moderate-severe groups reveal a high presence of cytokines, particularly interleukin-6 and the pro-inflammatory cytokines CCL2 and CCL7 (Supplementary Table 2.7). In contrast, proteins reduced in the moderate-severe group include CXCL5, which has been negatively correlated with these cytokines (54), and CCL17, whose decrease in severe patients has been previously published (55), (Supplementary Table 2.8). Among the metabolites of this same factor that are increased in moderate-severe groups, a substantial proportion are amino acids (17 of the top-50) (56, 57) and lipids (12 of the top-50), notably ceramides and other sphingolipids (58) (Supplementary Table 2.9). In contrast, the down-regulated metabolites include several phospholipids, lysophospholipids (59) and a range of xenobiotic compounds (Supplementary Table 2.10), which may reflect hepatic injury and impaired xenobiotic liver metabolism in COVID-19 (60).

Pathway-based multi-omics integration across disease severity was performed using the Pathway Integrative Analysis module. Comparisons were conducted in Multi-View mode for the groups mild vs. healthy, moderate vs. mild, and severe vs. moderate. The most pronounced separations were observed in the mild vs. healthy and moderate vs. mild comparisons, suggesting greater alterations in proteomic and metabolomic profiles during these stages (Figure 3G). All three comparisons revealed a high prevalence of categories related to immune response, particularly interleukin signaling (Supplementary Table 2.11). For instance, a visual exploration of the “Interleukin-4 and Interleukin-13 signaling” category (R-HSA-6785807) (Supplementary Table 2.12) shows a gradual increase in the levels of numerous cytokines from healthy to severe, with the most substantial increase occurring in the transition from mild to moderate (Figure 3H). The most relevant metabolomic categories in the mild vs. healthy and moderate vs. mild comparisons include nucleotide metabolism (Figure 3I, left panel) and amino acid and derivative metabolism (Figure 3I, right panel). Additionally, lipid metabolism exhibits significant alterations in both comparisons (Supplementary Table 2.13), aligning with the findings described above.

The reported study (46) analyzed the relationships between plasma proteins and metabolites using Spearman correlation and the construction of protein-metabolite regression models, with disease severity included as an interaction factor. This approach enabled the identification of significant protein-metabolite interactions and provided new insights into how plasma metabolic and proteomic profiles depend on disease severity. The use of TurbOmics complemented previous findings enabling the detection of key associations between proteins and metabolite classes linked to COVID-19 severity, particularly those involved in interleukin signaling and nucleotide, amino acid and lipid metabolism. Hence, our approach based on cutting-edge multivariate statistical techniques allowed projecting multi-omics data into a lower-dimensional space, efficiently capturing and highlighting the most significant sources of variation. This allowed us to replicate the identification of two major axes of variation in the healthy-mild and mild-moderate transitions originally described in the study. Furthermore, our method complemented the analysis by modeling the multi-omics data at the pathway level, integrating molecular signatures with knowledge base information and highlighting the pathways and functional categories that were altered in parallel with disease severity.

## DISCUSSION

Multi-omics integration provides a holistic view of biological systems by analyzing multiple functional layers simultaneously. Metabolomics is considered the omics level most closely linked to the phenotype, since it has the capacity to connect the genetic inheritance of an organism to the multiple environmental stimuli and characterize subsequent biological processes. Nevertheless, notable challenges still exist when analyzing metabolomics data in a multi-omics context. Here, we have introduced TurbOmics, a web-based platform for the analysis of metabolomics data using a multi-omics integrative approach. TurbOmics is a user-friendly web-based platform that enables users with different backgrounds to perform multi-omics analyses of untargeted and targeted metabolomics, transcriptomics and proteomics data. With an intuitive, accessible environment, TurbOmics does not require strong programming skills in applying state-of-the-art multivariate statistical techniques, enabling its use by researchers, technicians and physicians with different levels of experience. The platform provides interactive, exportable plots and tables that facilitate rapid and in-depth exploration of complex datasets in a multi-omics context.

The multi-omics integration performed by TurbOmics requires matching samples across omics layers, enabling direct identification of correlations and shared patterns among biomolecules based on the omics data itself (23). When omics data is obtained from the same biospecimens, the platform allows users to evaluate direct relationships between biomolecules across different functional levels. For omics data obtained from the same individual but different biospecimens (*e.g.*, plasma, tissue, urine), integrated analysis can corroborate findings across omics modalities or evaluate whether results in one layer can serve as a biomarker for changes occurring at another level (61). In addition, TurbOmics allows single-omics experiments to be analyzed, making it a valuable tool for a wide range of research contexts.

TurbOmics is unique in its specific support for untargeted metabolomics in multi-omics studies. Conversely to other omics, in untargeted metabolomics for the same feature multiple putative annotations could be generated, posing significant challenges for their handling and generally resulting in their exclusion from subsequent analyses. Yet, TurbOmics enables their handling by accepting user-obtained putative annotations from different tools, such as MetaboSearch, METLIN o CMM, and then process them with TurboPutative. Alternatively, the platform can automatically obtain the putative annotations by sequentially executing CMM and TurboPutative. This allows mitigating the identification bottleneck and facilitating data analysis in a multi-omics context. In addition, users can upload essential biochemical information, such as metabolite identification, biochemical class, spectral quality information, or identification confidence. This information is combined with multivariate analysis results, allowing a more comprehensive and in-depth interpretation of the untargeted metabolomics data. As showcased by our case studies, the application of TurbOmics allows the effective integration of both untargeted and targeted metabolomics within a multi-omics framework. In the first application, TurbOmics enabled the automated generation of putative annotations, facilitating the inclusion of all detected features in downstream quantitative analysis. Combined with the implemented multivariate statistical techniques, TurbOmics enabled a more in-depth multi-omics integration, uncovering a larger number of features with relevant abundance patterns. In contrast, the use of TurbOmics in the COVID-19 dataset highlights its ability to allow fast, intuitive and comprehensive data exploration, complementing the results of the original study by applying joined dimensionality reduction techniques and performing pathway-based multi-omics integration. This allows incorporating knowledge base information, thereby facilitating functional and biological interpretation. To the best of our knowledge, no other web-based platform addresses the challenges posed by putative annotations while enabling multi-omics integration with proteomics and transcriptomics data.

A further advantage of TurbOmics is the incorporation of state-of-the-art algorithms designed for multi-omics integration, providing a user-friendly interface with interactive capability for result exploration. Specifically, Multi-Omics Factor Analysis and Pathway Integrative Analysis modules are based on the MOFA and PathIntegrate frameworks, respectively. These algorithms employ dimensionality reduction approaches that effectively manage high-dimensional data with limited sample sizes, a known challenge in multi-omics studies. Leveraging these approaches, TurbOmics can efficiently compress high-dimensional data while preserving essential characteristics for further functional exploration, aiding researchers in extracting meaningful biological insights from complex multi-layered datasets without requiring coding and statistical skills.

Despite the significant advancements that TurbOmics offers for multi-omics studies, some limitations should be acknowledged. For instance, many biological studies generate omics data from different biological specimens or individuals, resulting in unpaired samples that cannot be integrated using TurbOmics. To mitigate this limitation, TurbOmics supports independent analyses of individual omics datasets, enabling users to benefit from the platform’s interactivity and visualizations while conducting statistical and functional analyses separately, and qualitatively comparing the results. Additionally, TurbOmics currently focuses on integrating metabolomics, proteomics, and transcriptomics, excluding other potentially relevant omics. Upcoming versions of TurbOmics aim to incorporate additional omics data types, such as genomics and epigenomics, thereby broadening its use in multi-omics research.

In summary, TurbOmics presents an accessible and innovative solution for multi-omics integration, specifically facilitating the analysis of metabolomics data in complex studies. Designed to support open science and collaborative research, TurbOmics is freely accessible without registration and generates exportable, standardized outputs that facilitate easy data sharing and integration within broader research frameworks. By supporting a transparent workflow and fostering multi-disciplinary collaborative research, TurbOmics has the potential to drive new discoveries in systems biology, making advanced multi-omics analysis available to the scientific community.

## MATERIAL AND METHODS

### Web server implementation

TurbOmics is hosted on a high-performance computing server running CentOS 7.3 Linux and is accessible via web browser. The back-end was developed using the Node.js framework, with core scripts for statistical analysis written in Python and R. The front-end was built using the the Next.js React framework along with several supporting libraries. TurbOmics has been tested on multiple web browsers, including Microsoft Edge, Google Chrome, and Mozilla Firefox, across major platforms (Windows, Linux, and macOS). Additionally, an official TurbOmics Docker image is available on the Docker Hub for local deployment of the web application.

TurbOmics is licensed under a Creative Commons Attribution-NonCommercial-NoDerivs 4.0 Unported License https://creativecommons.org/licenses/by-nc-nd/4.0/, and the source code is available on GitHub repository (see data availability).

### Data analysis implementation

#### Data preprocessing

Data preprocessing is performed on the back-end. Variance stabilization and normalization are executed using the *vsn* package (62) from Bioconductor project in R. ‘Center & Scale’ option applies feature-wise centering and scaling by subtracting the mean and scaling to unit variance. Subsequently, several algorithms can be selected by the user to perform missing value imputation. Both standardization and imputation are implemented using Python *scikit-learn* library. If k-nearest neighbors (KNN) method is used for imputation, the number of neighbors parameter is set to 3.

#### Putative Annotations

Putative annotations of metabolomics can be uploaded to the platform in the metabolomics metadata table. Alternatively, users can obtain these annotations through CEU Mass Mediator 3.0 (CMM) (32) and TurboPutative (22), both of which are run by TurbOmics. Specifically, CMM *batch* RESTful API is used to annotate features based on their m/z ratio using information from various external databases, including the Human Metabolome Database (63), KEGG (30), and LipidMaps (64). The CMM results are then processed by the four modules of TurboPutative: Tagger, REname, RowMerger, and TPMetrics. All tables generated during this process are accessible to the user; however, only the final table, TPFilter, will be used for subsequent multi-omics integration in the TurbOmics environment.

#### Exploratory Data Analysis

The distribution of abundance data is visualized using density curves and boxplots. Density curves are estimated from histograms normalized to have a total area of 1, with the number of bins determined by Sturges’ rule. Additionally, linear dimensionality reduction is performed using Principal Component Analysis (PCA) with Python library *scikit-learn*. Univariate linear regression between PCA scores and sample metadata is performed using the Python library *statsmodels*.

#### Multi-Omics Factor Analysis

The Python libraries *mofapy2* and *mofax* are used to run MOFA model (36). The algorithm is executed in an unbiased manner (no groups are specified) starting with an initial set of 30 factors; factors explaining less than 1.5% of the variation are eliminated, and the maximum number of iterations is set to 10, 000. All other parameters are kept at their default settings. Univariate linear regression between sample factor scores and sample metadata is performed using the Python *statsmodels* library. Over-Representation Analysis (ORA) with the top-n proteins and transcripts most associated with each factor is conducted using the g:GOSt RESTful API from g:Profiler (65), leveraging the GO (66), KEGG (30), and Reactome (31) knowledge bases. The entire set of proteins and transcripts uploaded by the user constitutes the domain scope used for enrichment. ORA analysis for metabolomics data is computed in the back-end using the Python library *SciPy* with the functional categories of KEGG and Reactome.

#### Pathway Integrative Analysis

The PathIntegrate modelling framework is employed to perform pathway-based multi-omics integration (37). In this framework, single-sample pathway analysis (ssPA) (67) is used to transform molecular-level abundance data into pathway level matrices. In our implementation, this transformation is performed using Singular Value Decomposition (SVD) (68) and Reactome database. TurbOmics incorporated both PathIntegrate supervised learning approaches: Single-View and Multi-View (see more in (37)). In our implementation, the Single-View approach employs Partial Least Squares Regression from *scikit-learn*, while Multi-View approach uses Multi-Block Partial Least Squares (MB-PLS) similar to PathIntegrate. Five latent variables are computed, and Reactome pathways containing more than three mapped biomolecules are included in the analysis. To assess model reliability, an empirical p-value is calculated over the R-squared value by performing permutations on the response variable. To estimate the contribution of each pathway to the model, variable importance in projection (VIP) is computed.

#### Enrichment Analysis

Functional class scoring enrichment analysis is performed using the GSEA algorithm (69), implemented in the R library *fgsea* (70) and utilizing the Hallmark (71), GO (66), KEGG (30), and Reactome (31) knowledge bases. Interactive enrichment analysis with user-specified categories through the web platform is run on the front-end, applying the GSEA algorithm implemented in JavaScript and calculating an empirical p-value obtained by performing 100 permutations. The Python *mummichog* library is used to perform the enrichment analysis of untargeted metabolomics data with the Mummichog model (72), setting the significance cutoff to the top 10% of the analyzed features.

### Datasets

#### Atherosclerosis data

Liver proteomics and metabolomics data were obtained from the original study, where full experimental procedures and pre-processing details are available (38). Metabolomics and proteomics datasets are accessible via the Metabolomics Workbench (Project ID: PR000985) at metabolomicsworkbench.org and PeptideAtlas at ftp://PASS01607:CP3575gq@ftp.peptideatlas.org/, respectively. RNA-seq data were collected from the same liver samples used in the original study. Approximately 50 mg of tissue were homogenized in TRIzol (Life Technologies, Madrid, Spain) and processed with chloroform. After centrifugation, RNA was purified using the RNeasy Mini Kit (Qiagen, Hilden, Germany). RNA concentration and integrity were assessed using a NanoDrop spectrophotometer (Thermo Fisher Scientific, Wilmington, DE, USA) and an Agilent 2100 Bioanalyzer (Agilent Technologies, Foster City, CA, USA). RNA sequencing was carried out by BGI Genomics company (Shenzhen, China). Briefly, mRNA was enriched from total RNA using oligo(dT) magnetic beads, fragmented, and reverse transcribed into cDNA. Libraries were prepared through end repair, adapter ligation, and PCR amplification. Single-stranded circular DNA libraries were generated and sequenced on the DNBSEQ platform (paired-end 150 bp, PE150).

Paired-end reads from DNBSEQ were processed using a pipeline that included filtering of low-quality reads with SOAPnuke (73) v1.5.6 and gene expression quantification from identified transcripts using RSEM (74) v1.3.1. RSEM internally aligned the reads to the mouse reference genome GRCm39.105 using bowtie2 (75) v2.4.5. Genes with a counts-per-million (CPM) value ≤1 in more than 50% of samples in each of the three mice groups were removed. The remaining 11, 999 genes were normalized using the TMM method (76) and log2-transformed for downstream multi-omics integration. TurbOmics analysis are accessible via the “Find Job” option on the web platform using the job ID “4eYOz8s8nD”.

#### COVID-19 data

Plasma proteomics and metabolomics data from the COVID-19 study are provided in Supplementary Tables S1.3 and S1.4 of the original study (46). Plasma protein concentrations were measured using the ProSeek Cardiovascular II, Inflammation, Metabolism, Immune Response, and Organ Damage panels (Olink Biosciences, Uppsala, Sweden), while plasma metabolomics data were acquired using the Global Metabolomics platform from Metabolon (Morrisville, NC, USA) via UHPLC-MS/MS. Full details of the multi-omics datasets and pre-processing are available in the original publication (46). TurbOmics analysis are accessible via the “Find Job” option on the web platform with the job ID “dtkgbe8mId”.

## DATA AVAILABILITY

TurbOmics is freely accessible without registration at https://proteomics.cnic.es/TurboPutative/TurbOmicsApp.html. A Docker image of TurbOmics is also available on Docker Hub at https://hub.docker.com/repository/docker/rbarreror/turboputative. The source code for the TurbOmics web server can be found on GitHub at https://github.com/CNIC-Proteomics/TurboPutative-web and https://github.com/CNIC-Proteomics/TurboOmics.

## SUPPLEMENTARY DATA

Supplementary Data are available at bioRxiv online.

## AUTHOR CONTRIBUTIONS

RB-R: Conceptualization, design, and programming of the web application, and manuscript writing. JMR, TN and WH: Supervision of design and methodology. MJ-F: RNA-seq analysis. AR: Supervision of the biological analysis. AM, AF, and JV: Supervision of the project and revision of the manuscript.

## ACKNOWLEDGEMENTS

We would like to thank to the members of the Cardiovascular Proteomics Laboratory and Proteomics Unit at CNIC, especially to Juan Antonio López and Estefanía Núñez for their invaluable feedback, which improved the design and fixing bugs. We are also thankful to the Quantitative Biology and Statistics Group at EMBL for their advice, and we extend our gratitude to Alberto Gil-de-la-Fuente for his support and assistance in incorporating CEU Mass Mediator into the application.

---

**Supplementary Figure S1.**
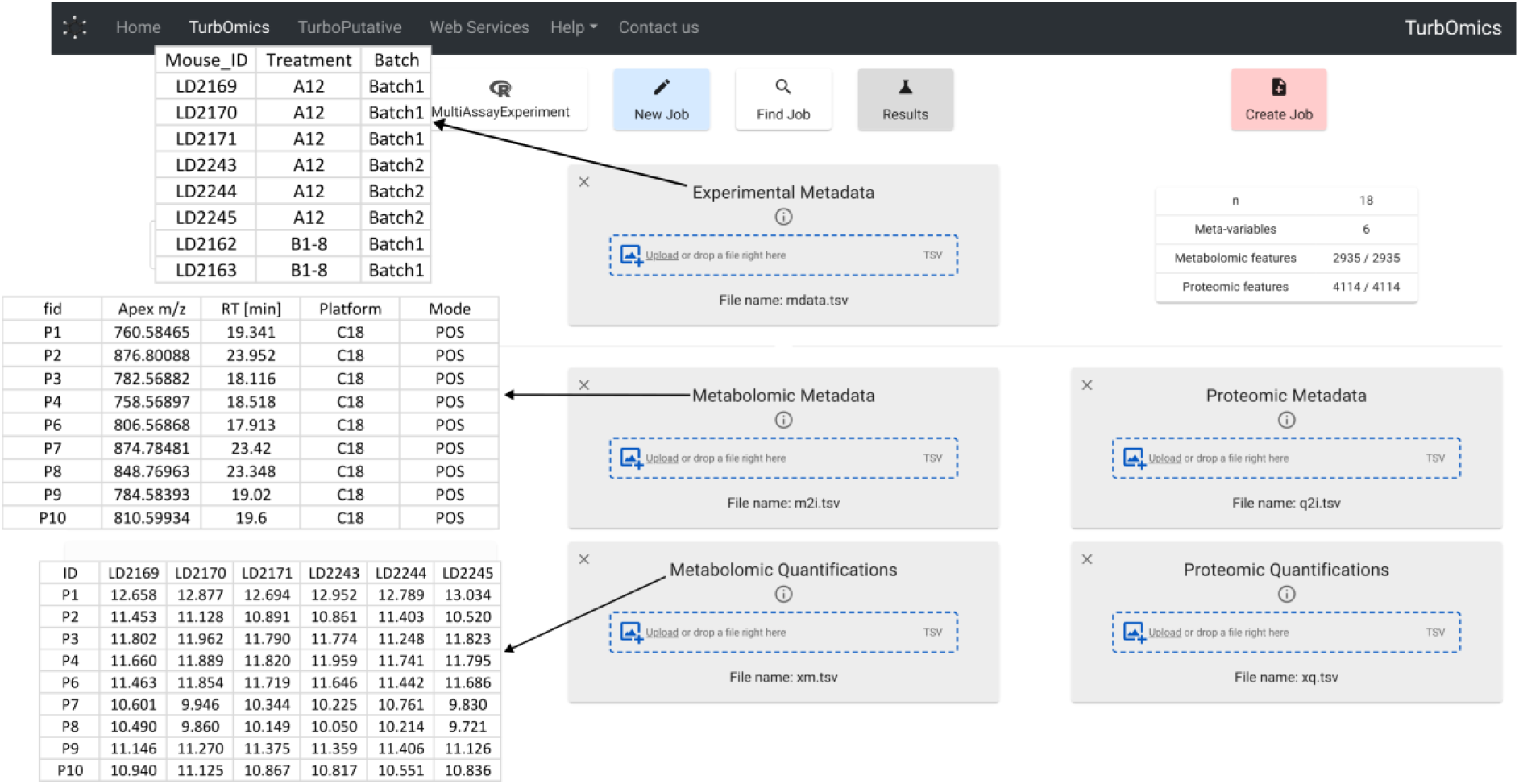
Data upload interface in the main page of TurbOmics.

**Supplementary Figure S2.**
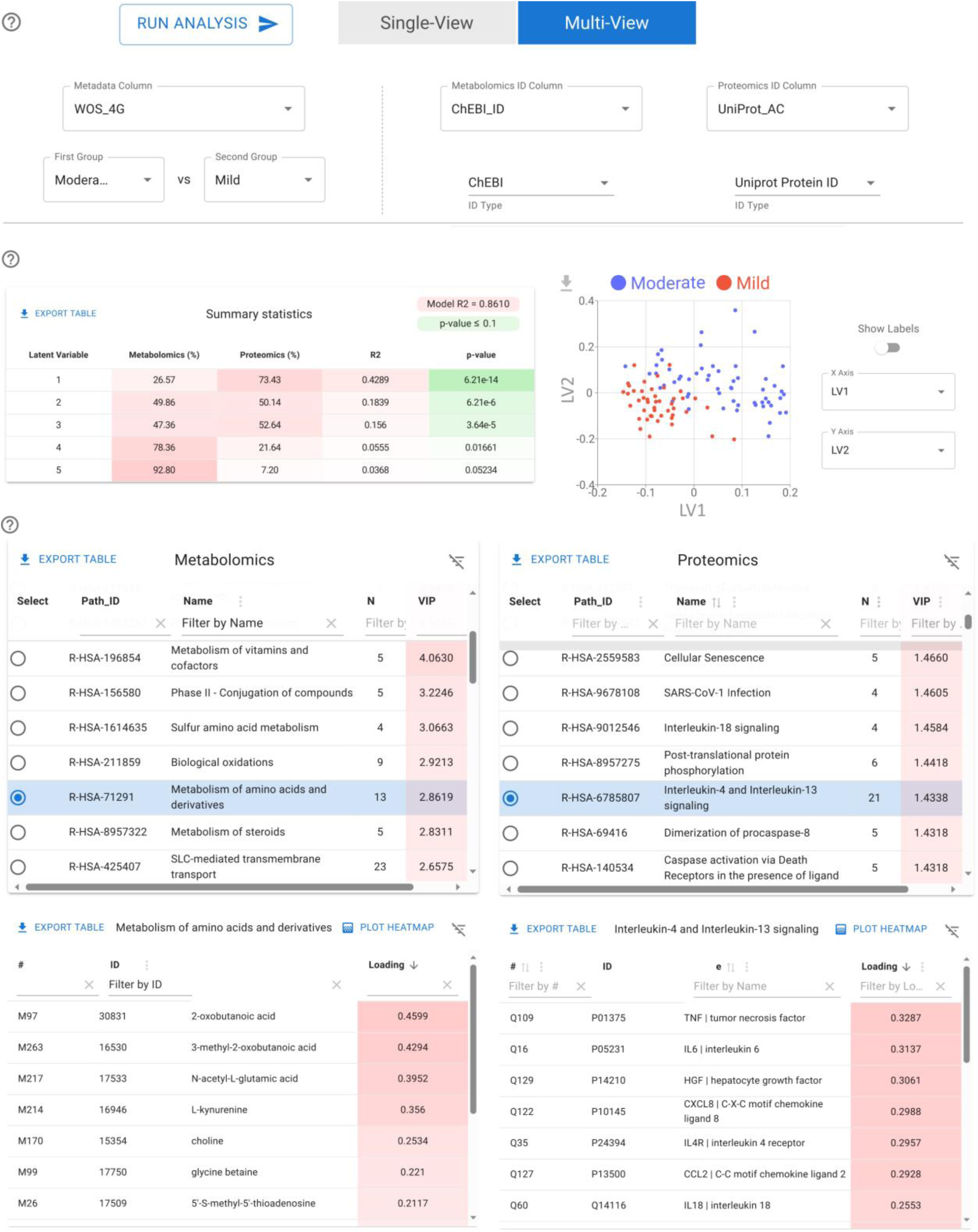
TurbOmics interface within the Pathway Integrative Analysis module executed in Multi-View mode.

**Supplementary Figure S3.**
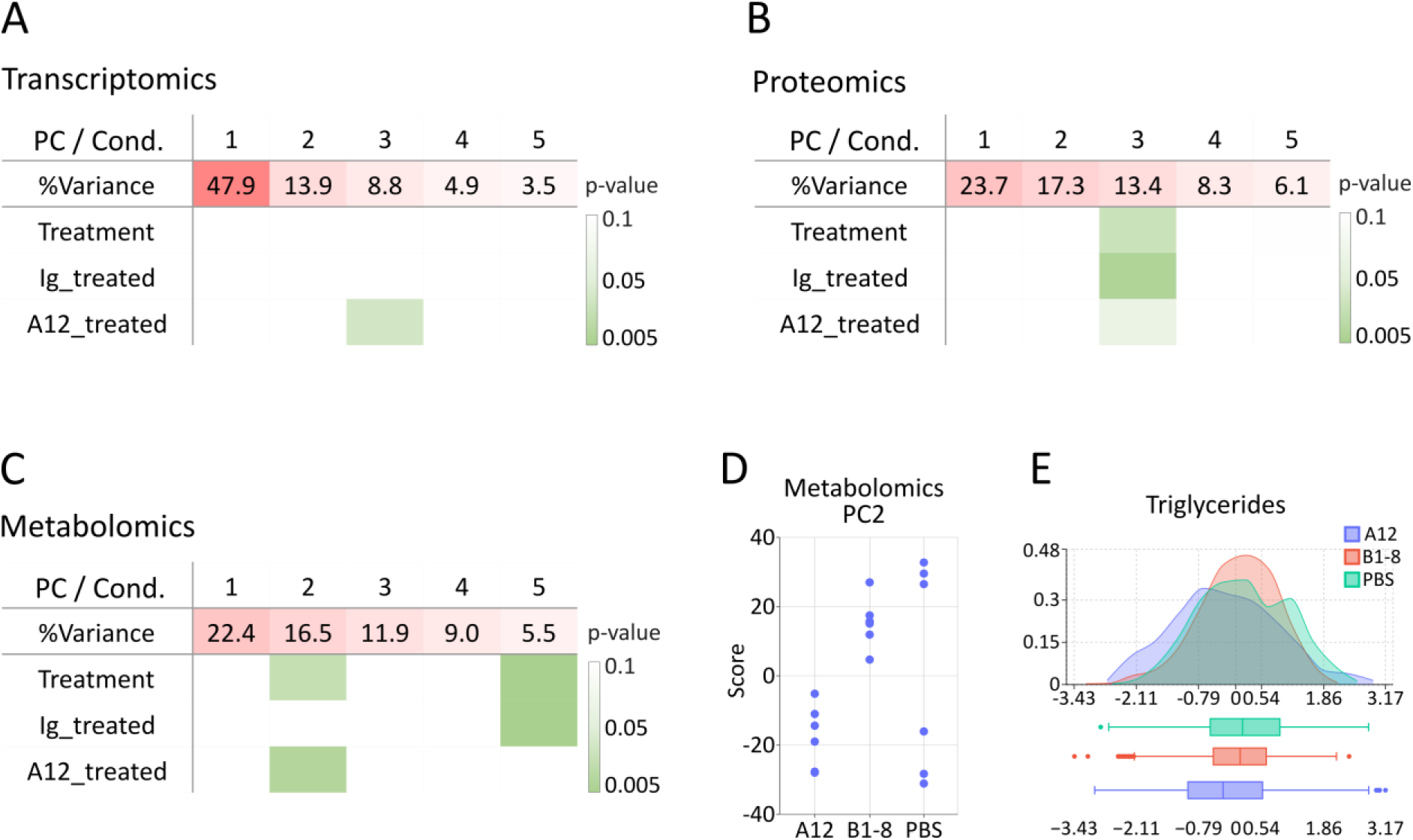
Exploratory data analysis in the atherosclerosis study using TurbOmics. (A), (B), and (C) present summary tables for the transcriptomics, proteomics, and metabolomics datasets, respectively, showing the principal components, their explained variability percentage, and the p-values for their association with treatment groups. Three group comparisons are presented: *Treatment* (A12 vs. B1-8 vs. PBS), *Ig_treated* (A12 and B1-8 vs. PBS), and *A12_treated* (A12 vs. B1-8 and PBS). (D) The scatter plot illustrates the distribution of scores for the second principal component across treatment groups. (E) Density and box plots depict the quantitative distribution of metabolomics features annotated as triglycerides, based on the Lipid Maps Structure Database.

## SUPPLEMENTARY TABLES LEGEND

Supplementary Tables S1. XLSX file containing tables S1.1–S1.7, with detailed information on the exploratory data analysis and multi-omics integration in the atherosclerosis study using TurbOmics.

Supplementary Tables S2. XLSX file containing tables S2.1–S2.13, with detailed information on the exploratory data analysis and multi-omics integration in the COVID-19 study using TurbOmics.

## REFERENCES

1. Mardis, E.R. (2008) Next-Generation DNA Sequencing Methods. Annual Review of Genomics and Human Genetics, 9, 387–402. 10.1146/annurev.genom.9.081307.164359

2. Nassar, A.F., Wu, T., Nassar, S.F. and Wisnewski, A.V. (2017) UPLC–MS for metabolomics: a giant step forward in support of pharmaceutical research. Drug Discovery Today, 22, 463– 470. 10.1016/j.drudis.2016.11.020

3. Guzman, U.H., Martinez-Val, A., Ye, Z., Damoc, E., Arrey, T.N., Pashkova, A., Renuse, S., Denisov, E., Petzoldt, J., Peterson, A.C., et al. (2024) Ultra-fast label-free quantification and comprehensive proteome coverage with narrow-window data-independent acquisition. Nature Biotechnology, 10.1038/s41587-023-02099-7. 10.1038/s41587-023-02099-7

4. Rodríguez-Lago, I., Blackwell, J., Mateos, B., Marigorta, U.M., Acosta, M.B. and Pollok, R. (2023) Recent Advances and Potential Multi-Omics Approaches in the Early Phases of Inflammatory Bowel Disease. Journal of Clinical Medicine, 12, 3418. 10.3390/jcm12103418

5. Rahimzadeh, N., Srinivasan, S.S., Zhang, J. and Swarup, V. (2024) Gene networks and systems biology in Alzheimer’s disease: Insights from multi-omics approaches. Alzheimer’s & Dementia, 20, 3587–3605. 10.1002/alz.13790

6. Hasin, Y., Seldin, M. and Lusis, A. (2017) Multi-omics approaches to disease. Genome Biology, 18, 83. 10.1186/s13059-017-1215-1

7. Czolk, R., Klueber, J., Sørensen, M., Wilmes, P., Codreanu-Morel, F., Skov, P.S., Hilger, C., Bindslev-Jensen, C., Ollert, M. and Kuehn, A. (2021) IgE-Mediated Peanut Allergy: Current and Novel Predictive Biomarkers for Clinical Phenotypes Using Multi-Omics Approaches. Frontiers in Immunology, 11. 10.3389/fimmu.2020.594350

8. Ivanisevic, T. and Sewduth, R.N. (2023) Multi-Omics Integration for the Design of Novel Therapies and the Identification of Novel Biomarkers. Proteomes, 11, 34. 10.3390/proteomes11040034

9. Xiao, Y., Bi, M., Guo, H. and Li, M. (2022) Multi-omics approaches for biomarker discovery in early ovarian cancer diagnosis. eBioMedicine, 79, 104001. 10.1016/j.ebiom.2022.104001

10. Morello, G., Salomone, S., D’Agata, V., Conforti, F.L. and Cavallaro, S. (2020) From Multi-Omics Approaches to Precision Medicine in Amyotrophic Lateral Sclerosis. Frontiers in Neuroscience, 14. 10.3389/fnins.2020.577755

11. Valenti, F., Falcone, I., Ungania, S., Desiderio, F., Giacomini, P., Bazzichetto, C., Conciatori, F., Gallo, E., Cognetti, F., Ciliberto, G., et al. (2021) Precision Medicine and Melanoma: Multi-Omics Approaches to Monitoring the Immunotherapy Response. International Journal of Molecular Sciences, 22, 3837. 10.3390/ijms22083837

12. Logotheti, M., Agioutantis, P., Katsaounou, P. and Loutrari, H. (2021) Microbiome Research and Multi-Omics Integration for Personalized Medicine in Asthma. Journal of Personalized Medicine, 11, 1299. 10.3390/jpm11121299

13. Eicher, T., Kinnebrew, G., Patt, A., Spencer, K., Ying, K., Ma, Q., Machiraju, R. and Mathé, E.A. (2020) Metabolomics and Multi-Omics Integration: A Survey of Computational Methods and Resources. Metabolites, 10, 202. 10.3390/metabo10050202

14. Pinu, F.R., Beale, D.J., Paten, A.M., Kouremenos, K., Swarup, S., Schirra, H.J. and Wishart, D. (2019) Systems Biology and Multi-Omics Integration: Viewpoints from the Metabolomics Research Community. Metabolites, 9, 76. 10.3390/metabo9040076

15. Fiehn, O. (2001) Combining genomics, metabolome analysis, and biochemical modelling to understand metabolic networks. Comparative and Functional Genomics, 2, 155–168. 10.1002/cfg.82

16. Nicholson, J.K., Lindon, J.C. and Holmes, E. (1999) ‘Metabonomics’: understanding the metabolic responses of living systems to pathophysiological stimuli via multivariate statistical analysis of biological NMR spectroscopic data. Xenobiotica, 29, 1181–1189. 10.1080/004982599238047

17. Roberts, L.D., Souza, A.L., Gerszten, R.E. and Clish, C.B. (2012) Targeted Metabolomics. Current Protocols in Molecular Biology, 98. 10.1002/0471142727.mb3002s98

18. Gowda, G.A.N. and Raftery, D. (2023) NMR Metabolomics Methods for Investigating Disease. Analytical chemistry, 95, 83. 10.1021/acs.analchem.2c04606 http://www.ncbi.nlm.nih.gov/pubmed/36625102

19. Pang, Z., Lu, Y., Zhou, G., Hui, F., Xu, L., Viau, C., Spigelman, A.F., MacDonald, P.E., Wishart, D.S., Li, S., et al. (2024) MetaboAnalyst 6.0: towards a unified platform for metabolomics data processing, analysis and interpretation. Nucleic Acids Research, 52, W398–W406. 10.1093/nar/gkae253

20. Tautenhahn, R., Patti, G.J., Rinehart, D. and Siuzdak, G. (2012) XCMS Online: A Web-Based Platform to Process Untargeted Metabolomic Data. Anal. Chem., 84, 5035–5039. 10.1021/ac300698c

21. Giacomoni, F., Le Corguillé, G., Monsoor, M., Landi, M., Pericard, P., Pétéra, M., Duperier, C., Tremblay-Franco, M., Martin, J.-F., Jacob, D., et al. (2015) Workflow4Metabolomics: a collaborative research infrastructure for computational metabolomics. Bioinformatics, 31, 1493–1495. 10.1093/bioinformatics/btu813

22. Barrero-Rodríguez, R., Rodriguez, J.M., Tarifa, R., Vázquez, J., Mastrangelo, A. and Ferrarini, A. (2022) TurboPutative: A web server for data handling and metabolite classification in untargeted metabolomics. Frontiers in Molecular Biosciences, 9. 10.3389/fmolb.2022.952149

23. Ewald, J.D., Zhou, G., Lu, Y., Kolic, J., Ellis, C., Johnson, J.D., Macdonald, P.E. and Xia, J. (2024) Web-based multi-omics integration using the Analyst software suite. Nature Protocols, 19, 1467–1497. 10.1038/s41596-023-00950-4

24. Liu, T., Salguero, P., Petek, M., Martinez-Mira, C., Balzano-Nogueira, L., Ramšak, Ž., McIntyre, L., Gruden, K., Tarazona, S. and Conesa, A. (2022) PaintOmics 4: new tools for the integrative analysis of multi-omics datasets supported by multiple pathway databases. Nucleic Acids Research, 50, W551–W559. 10.1093/nar/gkac352

25. Ding, J., Blencowe, M., Nghiem, T., Ha, S., Chen, Y.-W., Li, G. and Yang, X. (2021) Mergeomics 2.0: a web server for multi-omics data integration to elucidate disease networks and predict therapeutics. Nucleic Acids Research, 49, W375–W387. 10.1093/nar/gkab405

26. Zhou, G., Pang, Z., Lu, Y., Ewald, J. and Xia, J. (2022) OmicsNet 2.0: a web-based platform for multi-omics integration and network visual analytics. Nucleic Acids Research, 50, W527– W533. 10.1093/nar/gkac376

27. Kuo, T.-C., Tian, T.-F. and Tseng, Y.J. (2013) 3Omics: a web-based systems biology tool for analysis, integration and visualization of human transcriptomic, proteomic and metabolomic data. BMC Systems Biology, 7, 64. 10.1186/1752-0509-7-64

28. Zoppi, J., Guillaume, J.-F., Neunlist, M. and Chaffron, S. (2021) MiBiOmics: an interactive web application for multi-omics data exploration and integration. BMC Bioinformatics, 22, 6. 10.1186/s12859-020-03921-8

29. Zhou, G., Ewald, J. and Xia, J. (2021) OmicsAnalyst: a comprehensive web-based platform for visual analytics of multi-omics data. Nucleic Acids Research, 49, W476–W482. 10.1093/nar/gkab394

30. Kanehisa, M., Furumichi, M., Sato, Y., Kawashima, M. and Ishiguro-Watanabe, M. (2023) KEGG for taxonomy-based analysis of pathways and genomes. Nucleic Acids Research, 51, D587–D592. 10.1093/nar/gkac963

31. Milacic, M., Beavers, D., Conley, P., Gong, C., Gillespie, M., Griss, J., Haw, R., Jassal, B., Matthews, L., May, B., et al. (2024) The Reactome Pathway Knowledgebase 2024. Nucleic Acids Research, 52, D672–D678. 10.1093/nar/gkad1025

32. Gil-de-la-Fuente, A., Godzien, J., Saugar, S., Garcia-Carmona, R., Badran, H., Wishart, D.S., Barbas, C. and Otero, A. (2019) CEU Mass Mediator 3.0: A Metabolite Annotation Tool. Journal of Proteome Research, 18, 797–802. 10.1021/acs.jproteome.8b00720

33. MetaboSearch: Tool for Mass-Based Metabolite Identification Using Multiple Databases | PLOS ONE.

34. Montenegro-Burke, J.R., Guijas, C. and Siuzdak, G. (2020) METLIN: A Tandem Mass Spectral Library of Standards. Methods Mol Biol, 2104, 149–163. 10.1007/978-1-0716-0239-3_9 http://www.ncbi.nlm.nih.gov/pmc/articles/PMC10273217

35. Ramos, M., Schiffer, L., Re, A., Azhar, R., Basunia, A., Rodriguez, C., Chan, T., Chapman, P., Davis, S.R., Gomez-Cabrero, D., et al. (2017) Software for the Integration of Multiomics Experiments in Bioconductor. Cancer Research, 77, e39–e42. 10.1158/0008-5472.CAN-17-0344

36. Argelaguet, R., Velten, B., Arnol, D., Dietrich, S., Zenz, T., Marioni, J.C., Buettner, F., Huber, W. and Stegle, O. (2018) Multi-Omics Factor Analysis—a framework for unsupervised integration of multi-omics data sets. Molecular Systems Biology, 14. 10.15252/msb.20178124

37. Wieder, C., Cooke, J., Frainay, C., Poupin, N., Bowler, R., Jourdan, F., Kechris, K.J., Lai, R.P. and Ebbels, T. (2024) PathIntegrate: Multivariate modelling approaches for pathway-based multi-omics data integration. PLOS Computational Biology, 20, e1011814. 10.1371/journal.pcbi.1011814

38. Lorenzo, C., Delgado, P., Busse, C.E., Sanz-Bravo, A., Martos-Folgado, I., Bonzon-Kulichenko, E., Ferrarini, A., Gonzalez-Valdes, I.B., Mur, S.M., Roldán-Montero, R., et al. (2021) ALDH4A1 is an atherosclerosis auto-antigen targeted by protective antibodies. Nature, 589, 287–292. 10.1038/s41586-020-2993-2

39. Paapstel, K., Kals, J., Eha, J., Tootsi, K., Ottas, A., Piir, A., Jakobson, M., Lieberg, J. and Zilmer, M. (2018) Inverse relations of serum phosphatidylcholines and lysophosphatidylcholines with vascular damage and heart rate in patients with atherosclerosis. Nutrition, Metabolism and Cardiovascular Diseases, 28, 44–52. 10.1016/j.numecd.2017.07.011

40. Schmitz, G. and Ruebsaamen, K. (2010) Metabolism and atherogenic disease association of lysophosphatidylcholine. Atherosclerosis, 208, 10–18. 10.1016/j.atherosclerosis.2009.05.029

41. Moxon, J.V., Jones, R.E., Wong, G., Weir, J.M., Mellett, N.A., Kingwell, B.A., Meikle, P.J. and Golledge, J. (2017) Baseline serum phosphatidylcholine plasmalogen concentrations are inversely associated with incident myocardial infarction in patients with mixed peripheral artery disease presentations. Atherosclerosis, 263, 301–308. 10.1016/j.atherosclerosis.2017.06.925

42. Sutter, I., Klingenberg, R., Othman, A., Rohrer, L., Landmesser, U., Heg, D., Rodondi, N., Mach, F., Windecker, S., Matter, C.M., et al. (2016) Decreased phosphatidylcholine plasmalogens – A putative novel lipid signature in patients with stable coronary artery disease and acute myocardial infarction. Atherosclerosis, 246, 130–140. 10.1016/j.atherosclerosis.2016.01.003

43. Treede, I., Braun, A., Sparla, R., Kühnel, M., Giese, T., Turner, J.R., Anes, E., Kulaksiz, H., Füllekrug, J., Stremmel, W., et al. (2007) Anti-inflammatory Effects of Phosphatidylcholine *. Journal of Biological Chemistry, 282, 27155–27164. 10.1074/jbc.M704408200 http://www.ncbi.nlm.nih.gov/pubmed/17636253

44. Chen, M., Pan, Hongying, Dai, Yining, Zhang, Jiajie, Tong, Yongxi, Huang, Yicheng, Wang, Mingshan and and Huang, H. (2018) Phosphatidylcholine regulates NF-κB activation in attenuation of LPS-induced inflammation: evidence from in vitro study. Animal Cells and Systems, 22, 7–14. 10.1080/19768354.2017.1405072

45. Erős, G., Ibrahim, S., Siebert, N., Boros, M. and Vollmar, B. (2009) Oral phosphatidylcholine pretreatment alleviates the signs of experimental rheumatoid arthritis. Arthritis Research & Therapy, 11, R43. 10.1186/ar2651

46. Su, Y., Chen, D., Yuan, D., Lausted, C., Choi, J., Dai, C.L., Voillet, V., Duvvuri, V.R., Scherler, K., Troisch, P., et al. (2020) Multi-Omics Resolves a Sharp Disease-State Shift between Mild and Moderate COVID-19. Cell, 183, 1479–1495.e20. 10.1016/j.cell.2020.10.037

47. Tang, N., Yang, Y., Xie, Y., Yang, G., Wang, Q., Li, C., Liu, Z. and Huang, J. (2024) CD274 (PD-L1) negatively regulates M1 macrophage polarization in ALI/ARDS. Front. Immunol., 15, 1344805. 10.3389/fimmu.2024.1344805

48. Marlin, R., Godot, V., Cardinaud, S., Galhaut, M., Coleon, S., Zurawski, S., Dereuddre-Bosquet, N., Cavarelli, M., Gallouët, A.-S., Maisonnasse, P., et al. (2021) Targeting SARS-CoV-2 receptor-binding domain to cells expressing CD40 improves protection to infection in convalescent macaques. Nat Commun, 12, 5215. 10.1038/s41467-021-25382-0

49. Haljasmägi, L., Salumets, A., Rumm, A.P., Jürgenson, M., Krassohhina, E., Remm, A., Sein, H., Kareinen, L., Vapalahti, O., Sironen, T., et al. (2020) Longitudinal proteomic profiling reveals increased early inflammation and sustained apoptosis proteins in severe COVID-19. Sci Rep, 10, 20533. 10.1038/s41598-020-77525-w

50. Gudowska-Sawczuk, M. and Mroczko, B. (2022) The Role of Nuclear Factor Kappa B (NF-κB) in Development and Treatment of COVID-19: Review. Int J Mol Sci, 23, 5283. 10.3390/ijms23095283 http://www.ncbi.nlm.nih.gov/pmc/articles/PMC9101079

51. Nilsson-Payant, B.E., Uhl, S., Grimont, A., Doane, A.S., Cohen, P., Patel, R.S., Higgins, C.A., Acklin, J.A., Bram, Y., Chandar, V., et al. (2021) The NF-κB Transcriptional Footprint Is Essential for SARS-CoV-2 Replication. J Virol, 95, e0125721. 10.1128/JVI.01257-21 http://www.ncbi.nlm.nih.gov/pmc/articles/PMC8577386

52. Toro, D.M., da Silva-Neto, P.V., de Carvalho, J.C.S., Fuzo, C.A., Pérez, M.M., Pimentel, V.E., Fraga-Silva, T.F.C., Oliveira, C.N.S., Caruso, G.R., Vilela, A.F.L., et al. (2023) Plasma Sphingomyelin Disturbances: Unveiling Its Dual Role as a Crucial Immunopathological Factor and a Severity Prognostic Biomarker in COVID-19. Cells, 12, 1938. 10.3390/cells12151938

53. Marfia, G., Navone, S., Guarnaccia, L., Campanella, R., Mondoni, M., Locatelli, M., Barassi, A., Fontana, L., Palumbo, F., Garzia, E., et al. (2021) Decreased serum level of sphingosine-1-phosphate: a novel predictor of clinical severity in COVID-19. EMBO Mol Med, 13, e13424. 10.15252/emmm.202013424

54. Zaid, Y., Doré, É., Dubuc, I., Archambault, A.-S., Flamand, O., Laviolette, M., Flamand, N., Boilard, É. and Flamand, L. (2021) Chemokines and eicosanoids fuel the hyperinflammation within the lungs of patients with severe COVID-19. Journal of Allergy and Clinical Immunology, 148, 368–380.e3. 10.1016/j.jaci.2021.05.032

55. Sugiyama, M., Kinoshita, N., Ide, S., Nomoto, H., Nakamoto, T., Saito, S., Ishikane, M., Kutsuna, S., Hayakawa, K., Hashimoto, M., et al. (2021) Serum CCL17 level becomes a predictive marker to distinguish between mild/moderate and severe/critical disease in patients with COVID-19. Gene, 766, 145145. 10.1016/j.gene.2020.145145

56. Masoodi, M., Peschka, M., Schmiedel, S., Haddad, M., Frye, M., Maas, C., Lohse, A., Huber, S., Kirchhof, P., Nofer, J.-R., et al. (2022) Disturbed lipid and amino acid metabolisms in COVID-19 patients. J Mol Med, 100, 555–568. 10.1007/s00109-022-02177-4

57. Atila, A., Alay, H., Yaman, M.E., Akman, T.C., Cadirci, E., Bayrak, B., Celik, S., Atila, N.E., Yaganoglu, A.M., Kadioglu, Y., et al. (2021) The serum amino acid profile in COVID-19. Amino Acids, 53, 1569–1588. 10.1007/s00726-021-03081-w

58. Khodadoust, M.M. (2021) Inferring a causal relationship between ceramide levels and COVID-19 respiratory distress. Sci Rep, 11, 20866. 10.1038/s41598-021-00286-7

59. Wei, J., Liu, X., Xiao, W., Lu, J., Guan, L., Fang, Z., Chen, J., Sun, B., Cai, Z., Sun, X., et al. (2023) Phospholipid remodeling and its derivatives are associated with COVID-19 severity. Journal of Allergy and Clinical Immunology, 151, 1259–1268. 10.1016/j.jaci.2022.11.032

60. Hammoudeh, S.M., Hammoudeh, A.M., Bhamidimarri, P.M., Mahboub, B., Halwani, R., Hamid, Q., Rahmani, M. and Hamoudi, R. (2021) Insight into molecular mechanisms underlying hepatic dysfunction in severe COVID-19 patients using systems biology. World journal of gastroenterology, 27, 2850–2870. 10.3748/wjg.v27.i21.2850 http://www.ncbi.nlm.nih.gov/pubmed/34135558

61. Chu, S., Huang, M., Kelly, R., Benedetti, E., Siddiqui, J., Zeleznik, O., Pereira, A., Herrington, D., Wheelock, C., Krumsiek, J., et al. (2019) Integration of Metabolomic and Other Omics Data in Population-Based Study Designs: An Epidemiological Perspective. Metabolites, 9, 117. 10.3390/metabo9060117

62. Huber, W., Heydebreck, A. von, Sültmann, H., Poustka, A. and Vingron, M. (2002) Variance stabilization applied to microarray data calibration and to the quantification of differential expression. Bioinformatics, 18, S96–S104. 10.1093/bioinformatics/18.suppl_1.S96

63. Wishart, D.S., Guo, A., Oler, E., Wang, F., Anjum, A., Peters, H., Dizon, R., Sayeeda, Z., Tian, S., Lee, B.L., et al. (2022) HMDB 5.0: the Human Metabolome Database for 2022. Nucleic Acids Research, 50, D622–D631. 10.1093/nar/gkab1062

64. Conroy, M.J., Andrews, R.M., Andrews, S., Cockayne, L., Dennis, E.A., Fahy, E., Gaud, C., Griffiths, W.J., Jukes, G., Kolchin, M., et al. (2024) LIPID MAPS: update to databases and tools for the lipidomics community. Nucleic Acids Research, 52, D1677–D1682. 10.1093/nar/gkad896

65. Raudvere, U., Kolberg, L., Kuzmin, I., Arak, T., Adler, P., Peterson, H. and Vilo, J. (2019) g:Profiler: a web server for functional enrichment analysis and conversions of gene lists (2019 update). Nucleic Acids Research, 47, W191–W198. 10.1093/nar/gkz369

66. Aleksander, S.A., Balhoff, J., Carbon, S., Cherry, J.M., Drabkin, H.J., Ebert, D., Feuermann, M., Gaudet, P., Harris, N.L., Hill, D.P., et al. (2023) The Gene Ontology knowledgebase in 2023. GENETICS, 224. 10.1093/genetics/iyad031

67. Wieder, C., Lai, R.P.J. and Ebbels, T.M.D. (2022) Single sample pathway analysis in metabolomics: performance evaluation and application. BMC Bioinformatics, 23, 481. 10.1186/s12859-022-05005-1

68. Tomfohr, J., Lu, J. and Kepler, T.B. (2005) Pathway level analysis of gene expression using singular value decomposition. BMC Bioinformatics, 6, 225. 10.1186/1471-2105-6-225 http://www.ncbi.nlm.nih.gov/pubmed/16156896

69. Subramanian, A., Tamayo, P., Mootha, V.K., Mukherjee, S., Ebert, B.L., Gillette, M.A., Paulovich, A., Pomeroy, S.L., Golub, T.R., Lander, E.S., et al. (2005) Gene set enrichment analysis: A knowledge-based approach for interpreting genome-wide expression profiles. Proceedings of the National Academy of Sciences, 102, 15545–15550. 10.1073/pnas.0506580102

70. Sergushichev, A.A. (2016) An algorithm for fast preranked gene set enrichment analysis using cumulative statistic calculation. biorXiv, 10.1101/060012. 10.1101/060012

71. Liberzon, A., Birger, C., Thorvaldsdóttir, H., Ghandi, M., Mesirov, J.P. and Tamayo, P. (2015) The Molecular Signatures Database Hallmark Gene Set Collection. Cell Systems, 1, 417–425. 10.1016/j.cels.2015.12.004

72. Li, S., Park, Y., Duraisingham, S., Strobel, F.H., Khan, N., Soltow, Q.A., Jones, D.P. and Pulendran, B. (2013) Predicting Network Activity from High Throughput Metabolomics. PLoS Computational Biology, 9, e1003123. 10.1371/journal.pcbi.1003123

73. Chen, Y., Chen, Y., Shi, C., Huang, Z., Zhang, Y., Li, S., Li, Y., Ye, J., Yu, C., Li, Z., et al. (2018) SOAPnuke: a MapReduce acceleration-supported software for integrated quality control and preprocessing of high-throughput sequencing data. GigaScience, 7. 10.1093/gigascience/gix120

74. Li, B. and Dewey, C.N. (2011) RSEM: accurate transcript quantification from RNA-Seq data with or without a reference genome. BMC Bioinformatics, 12, 323. 10.1186/1471-2105-12-323

75. Langmead, B. and Salzberg, S.L. (2012) Fast gapped-read alignment with Bowtie 2. Nat Methods, 9, 357–359. 10.1038/nmeth.1923

76. Robinson, M.D. and Oshlack, A. (2010) A scaling normalization method for differential expression analysis of RNA-seq data. Genome Biology, 11, R25. 10.1186/gb-2010-11-3-r25

